# Local extirpation is pervasive among populations of Galápagos endemic tomatoes

**DOI:** 10.1101/814160

**Authors:** Matthew J.S. Gibson, Maria de Lourdes Torres, Leonie C. Moyle

## Abstract

The Galápagos Islands are home to incredible endemic biodiversity that is of high conservation interest. Two such endemic species are the Galápagos tomatoes: *Solanum cheesmaniae* and *Solanum galapagense*. Both are known from historical location records, but like many endemic plant species on the Galápagos, their current conservation status is unclear. We revisited previously documented sites of endemic species on San Cristóbal, Santa Cruz, and Isabela, and document the disappearance of >80% of these populations. In contrast, we find that two invasive relatives (*Solanum pimpinellifolium* and *Solanum lycopersicum*) are now highly abundant, and in some cases—based on morphological observations—might be hybridizing with endemics. Our findings suggest that expanding human developments and putative interspecific hybridization are among the major factors affecting the prevalence of invasives and the threatened persistence of the endemic populations.

## 1 Introduction

The Galápagos Islands are a hotspot for species endemism and a key destination for scientists and tourists alike. Nonetheless, recent human colonization (no earlier than 1535; Quiroga, 2018) and a boom in tourism over the past 50 years have amplified many threats to endemic plant and animal species. Despite extensive conservation efforts–especially focused on the islands’ charismatic megafauna–many species remain understudied in terms of their conservation status. The Galápagos tomatoes—*Solanum cheesmaniae* (L. Riley) Fosberg (CHS) and *Solanum galapagense* S.C. Darwin & Peralta (GAL)—are two such endemic species which are currently designated as least concern by the IUCN (based on collection data from the 1950s and 60s; Tye & Siemens, 2014), but which observational reports suggest may be at risk of local population extinction (Darwin, 2009; Nuez et al., 2004). Resolving this discrepancy requires current and systematic demographic data to assess the magnitude of potential conservation threats to these species.

Island plants commonly face three threats: displacement due to expanding urban developments, ecological and demographic displacement by invasive species, and loss of genetic integrity via interspecific hybridization (D’Antonio & Dudley, 1995; Francisco-Ortega et al., 2000; Olson, 1989; Raven, 1998). Insular ecosystems such as the Galápagos Islands are vulnerable to invasion due to low levels of species richness and reduced competition (Williamson, 1996). Furthermore, the detrimental effects of invasion often act synergistically with other anthropogenic and non-anthropogenic forces. For example, expanding human developments can facilitate the invasion of exotics via increased propagule pressure and increased habitat disturbance. Similarly, ecological displacement by invasives can be accompanied by genetic disruption via hybridization, when invasive and native species are close evolutionary relatives. Although the outcomes of hybridization are potentially numerous, the focus of conservation biologists is on its potential to drive rare endemic taxa to extinction, as both low and high relative hybrid fitness can potentially lead to the exclusion of parental populations (Todesco et al., 2016; Wolf et al., 2001) or to the extinction of purely endemic genomes (Muhlfeld et al., 2014; Todesco et al., 2016).

Prior data suggest that the Galápagos tomatoes might be exemplars of these three aforementioned threats. Economic, touristic, and agricultural development on the islands have been explosive over the past 60 years (Quiroga, 2018). The accompanying economic transition from primarily agrarian to tourism-focused (Pizzitutti et al., 2017) and the subsequent expansion of coastal towns has affected natural habitats as well as provided new habitats for invaders (e.g. in abandoned highland crop fields, industrial sites, solid waste plants, and along roadsides (Darwin, 2009; Quiroga, 2018). Among these invasive species are *Solanum pimpinellifolium* L. (PIM; currant tomato) and *Solanum lycopersicum* L. (LYC; domesticated tomato; Darwin, 2003; Rick, 1963) which are thought to have been introduced to the islands after human colonization (Darwin et al., 2003; Darwin, 2009) and partially overlap with the endemics in terms of their geographic distributions (Darwin, 2009; Nuez et al., 2004). Where sympatric, these invasive species represent potential demographic and genetic threats. Unlike their invasive relatives, both GAL and CHS have extremely high seed dormancy (less than 1% germination without bleach treatment; Rick, 1956; Rick & Bowman, 1961), amplifying the opportunity for their local competitive exclusion in sympatry. In addition, hybrids between invasives and endemics have been reported (Darwin, 2009). Since hybrids between CHS and LYC do not display the low germination rates seen in the endemic taxa (Rick, 1956), and could thereby have a competitive advantage in the native parental environments, this hybridization could threaten endemics both directly via reproductive loss and long-term via competitive exclusion.

The presence of these multiple simultaneous conservation threats suggests that there is a high potential for displacement and/or extinction of populations of the two endemic tomato species. Consistent with this, field work by Darwin (2009) and Nuez et al. (2004) in 2000 and 2002 suggested that several previously documented endemic populations have disappeared. Using collection data for 87 accessions maintained by the C.M. Rick Tomato Genetics Resource Center (TGRC, Davis, Calif., USA; http://tgrc.davis.edu) and those documented by Darwin (2009) and Nuez et al. (2004), we visited historical population sites on 3 of the 13 major Galápagos islands. We report quantitative field data from 2018 and 2019 that indicate further loss of endemic populations of CHS and GAL. We also found that invasive species are highly abundant relative to endemics in the areas surveyed. Furthermore, we document the presence of populations with intermediate morphologies, suggesting the possibility of past or ongoing invasive-endemic and endemic-endemic hybridization.

## 2 Methods

We visited San Cristóbal (2018 & 2019), Santa Cruz (2018 & 2019), and Isabela (2018)–the three largest and most populated islands in the archipelago. These islands contain 40 of the 87 documented TGRC locations (TGRC passport data) and these sites, as well as those described in Darwin (2009) and Nuez et al. (2004), were given priority during our expeditions. We searched for plants (or remnants of plants) within a 200m radius of each revisited collection site. Additionally, focusing on areas ecologically similar to documented collection sites and/or previously observed to contain tomato plants by national park rangers, we searched for additional undocumented populations. Population identity was determined based on the taxonomic treatments in Darwin et al. (2003), with the exception of LYC, for which we distinguish two forms—LYC (domesticated tomato) and LYC var. cerasiforme (CER; cherry tomato; Rick, 1956)—to maintain consistency with Nuez et al’s (2004) previous description of several populations of the latter. CER has been described on the Galápagos as having small and irregularly shaped oblate fruits and typically lacking trichomes (Rick, 1956). Ripe fruit color was the major diagnostic trait used in species identification, as endemic fruit color varies from orange to yellow while invasives are usually deep red. We also measured nine additional traits (listed in Table 2 & Table 3) previously identified as variable, and diagnostic, among island taxa in Darwin (2003). Populations were sampled across linear transects and detailed photographs of leaf, fruit, and flower morphology were taken of 10-15 individuals per population against color and length standards. Leaf morphology and fruit color (RGB and HSV) were measured on select populations using *imageJ* (Schindelin et al., 2012). The latitude and longitude of each collection location was logged using a smartphone. Each site and the identity of species found within are described in Table 1 and displayed in Figure 1.

**Table 1:**
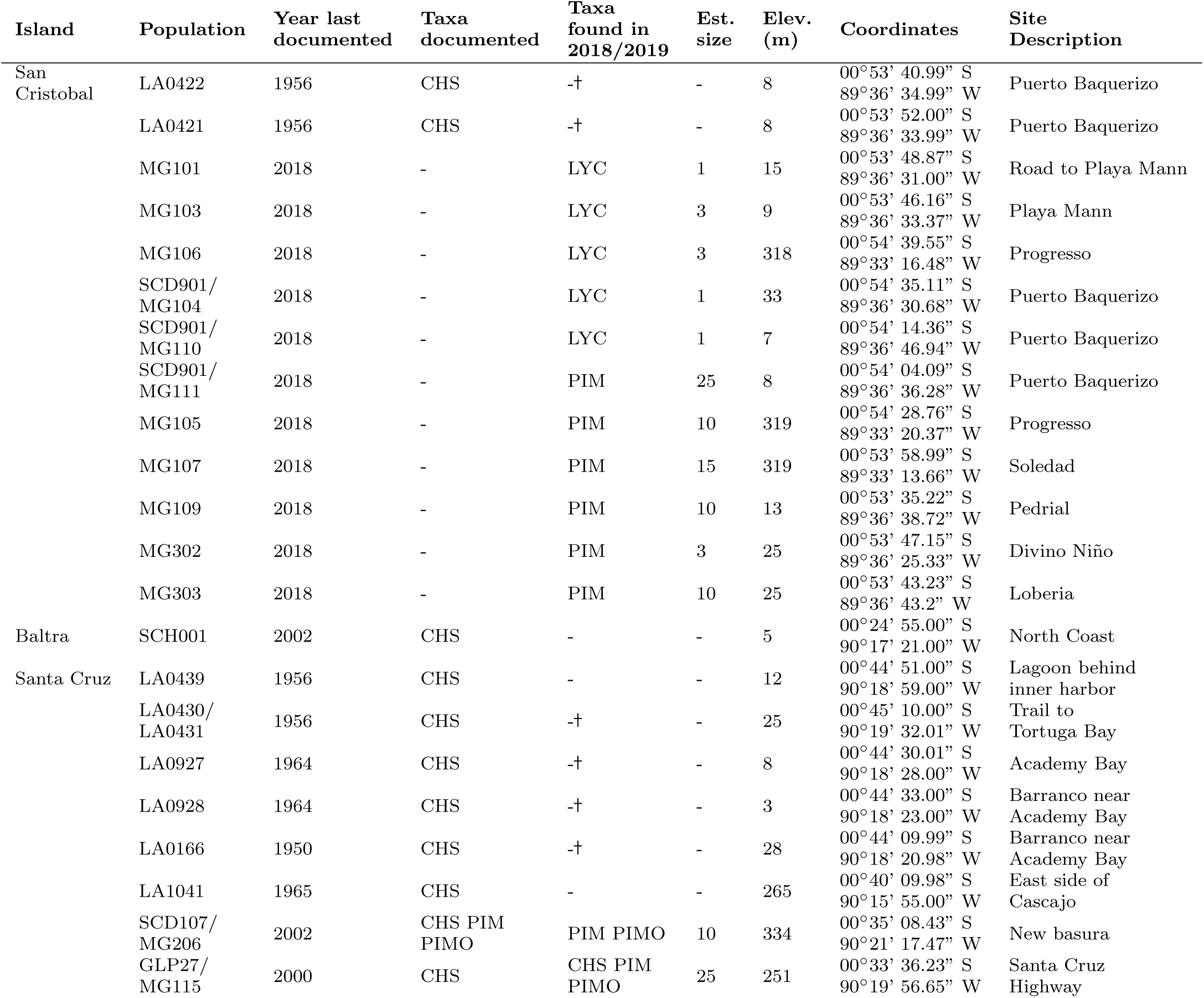
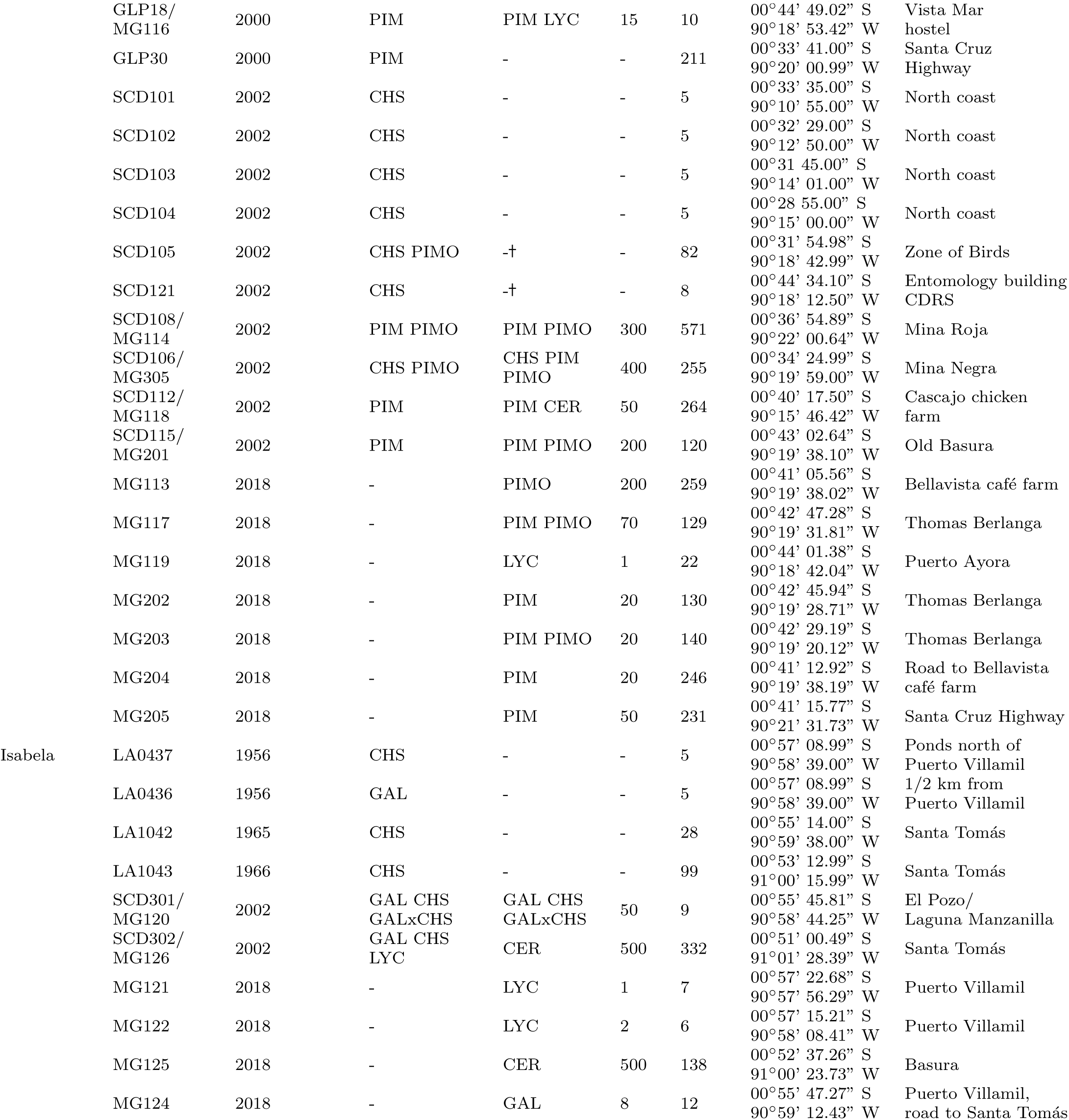
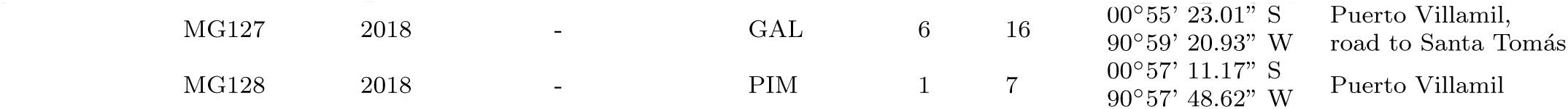
List of collections sites, both historical and new, visited during the May 2018 and June 2019 field seasons. Species originally reported to be at historical collection sites as well as the species observed at these and new sites in 2018/2019 are shown in adjacent columns. Species codes are as follows: (CHS) *S. cheesmaniae*; (GAL) *S. galapagense*; (LYC) *S. lycopersicum*; (PIM) *S. pimpinellifolium*; (PIMP) Orange-fruited *S. pimpinellifolium*; (GALxCHS) S. cheesmaniae X S. galapagense hybrid. Historical populations that were classified as *S. cheesmaniae* X *S. pimpinellifolium* hybrids by Darwin (2009) are represented here as orange-fruited *S. pimpinellifolium* for simplicity. † Population missing across consecutive years

**Table 2.**
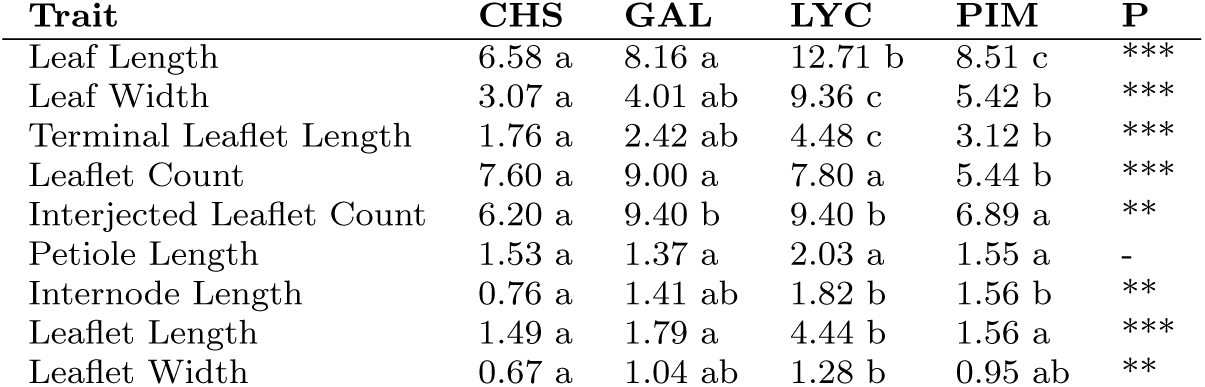
ANOVA tests of morphological differences between species. (* P < 0.05; ** P < 0.01; *** P < 0.001). No putative hybrids were included in these tests.

**Table 3.** Morphological comparison of orange and red-fruited PIM morphs. Data from population MG114. (* P < 0.05; ** P < 0.01). With the exception of color, plants are phenotypically identical.

| Trait | Orange<br>PIM Mean | Red<br>PIM Mean | Difference<br>(95% CI) |
| --- | --- | --- | --- |
| Leaf Length | 8.33 | 8.66 | -0.32 (-3.41, 2.77) |
| Leaf Width | 5.34 | 5.49 | -0.15 (-2.01, 1.71) |
| Terminal Leaflet Length | 3.11 | 3.22 | -0.11 (-1.26, 1.04) |
| Leaflet Count | 5.5 | 5.4 | 0.1 (-1.46, 1.66) |
| Interjected Leaflet Count | 5.75 | 7.8 | -2.05 (-3.65, -0.45)* |
| Petiole Length | 1.42 | 1.66 | -0.24 (-0.84, 0.37) |
| Internode Length | 1.58 | 1.55 | 0.04 (-0.53, 0.60) |
| Leaflet Length | 2.69 | 2.47 | 0.22 (-0.88, 1.32) |
| Leaflet Width | 1.03 | 0.88 | 0.15 (-0.58, 0.88) |
| Color PC1 | -1.96 | 1.57 | -3.53 (-4.82, -2.24)** |

**Fig. 1.**
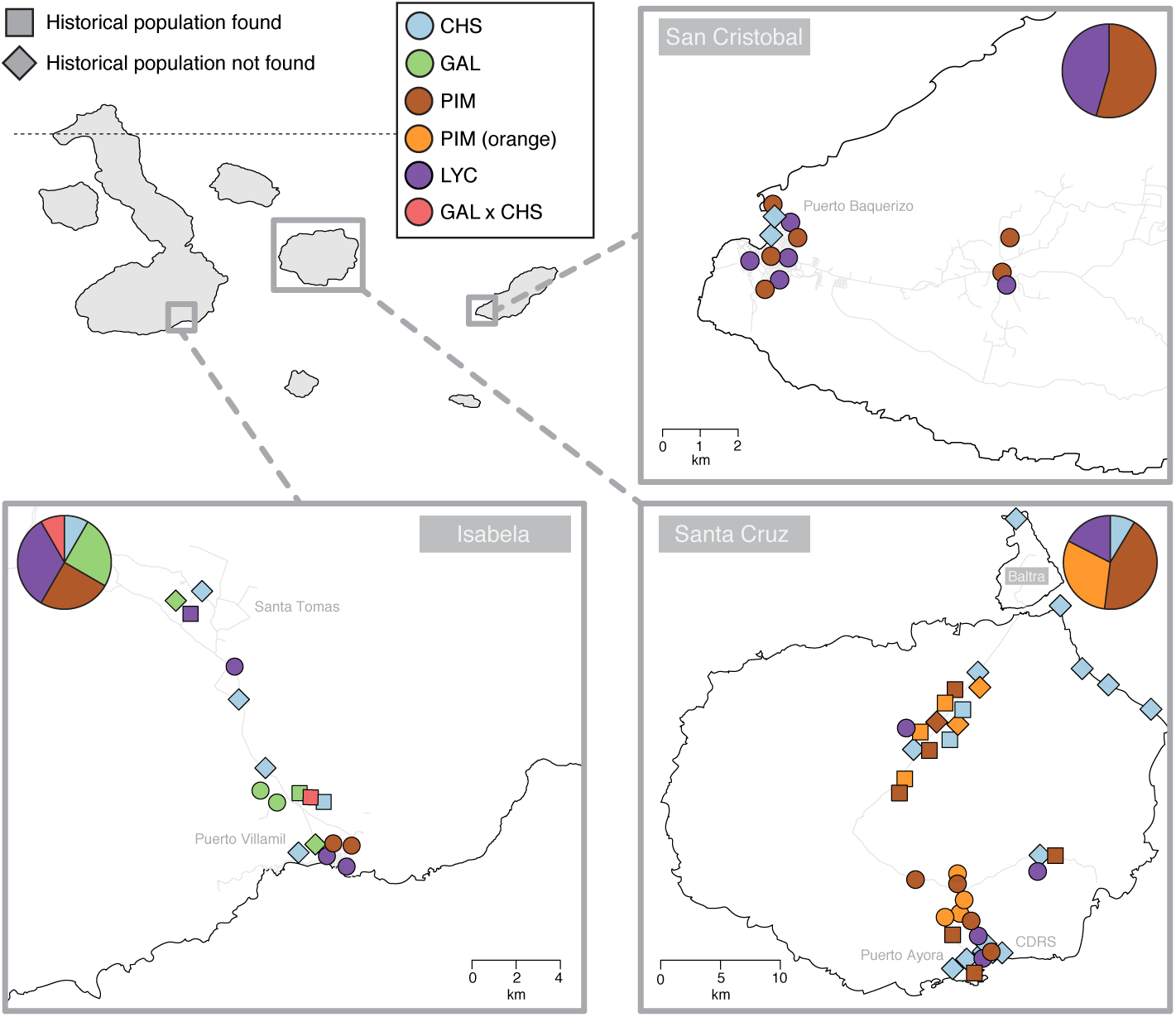
Maps of collection sites on Isabela, Santa Cruz, and San Cristóbal. Diamonds represent historical populations that were not relocated during this survey. Squares represent historical populations that were relocated. Circles represent newly discovered populations. Pie-charts show overall species abundance on each island. Most historical populations were not found in the contemporary survey. CER populations are represented as LYC for simplicity.

To evaluate evidence for trait differences between groups (species or populations), we performed analysis of variance (ANOVA) and post-hoc Tukey tests. Principle components analysis (PCA) was also used to summarize total phenotypic variation into two major explanatory axes. All statistical analyses were performed using R ver. 3.5.1 (R Core Team, 2019). Post-hoc tests were implemented using the R package *agricolae* (Mendiburu, 2019).

## 3 Results

Across the three islands, we visited 24 sites documented to contain endemic populations of CHS or GAL. Only four populations could be located after exhaustive searching: one CHS population on Santa Cruz Highway, Santa Cruz (N ≈ 20), one CHS population at El mina Granillo Negro (N > 300), and one site containing both CHS and GAL at the well (El Pozo) on Isabela (N ≈ 50). Six of the missing populations—whose presence was initially documented by Darwin (2009) or Nuez et al. (2004)—have been lost within only the past 16 years. In contrast to the rarity of endemics, we documented 30 sites containing invasive individuals of PIM, LYC, CER, or both PIM and LYC, across the three islands, with LYC being the predominant species on Isabela and PIM being the predominant species on San Cristóbal and Santa Cruz (Table 1; Figure 1). Population sizes at these sites varied from 1 to approximately 500 individuals (Table 1). Of these 30 locations, only seven had been previously reported as invasive populations (Darwin, 2009; Nuez et al., 2004).

### 3.1 Morphology

Consistent with the taxonomic treatments described in Darwin (2003), we detect significant morphological differentiation of the four Galápagos taxa (Table 2; Figure 2). ANOVA tests for eight of nine diagnostic trait differences between species were statistically significant (P < 0.01; Table 2), and the first two axes of a principle components analysis clearly cluster each taxon (Figure 2A). The first PC axis (R^2^ = 27.76%) mostly separates invasive from endemic species, while the second axis (R^2^ = 17.45%) differentiates between CHS and GAL. GAL collections from site MG120 are remarkably invariant (Green dots in Figure 2A), whereas CHS, PIM, and LYC are morphologically much more diverse.

**Fig. 2.**
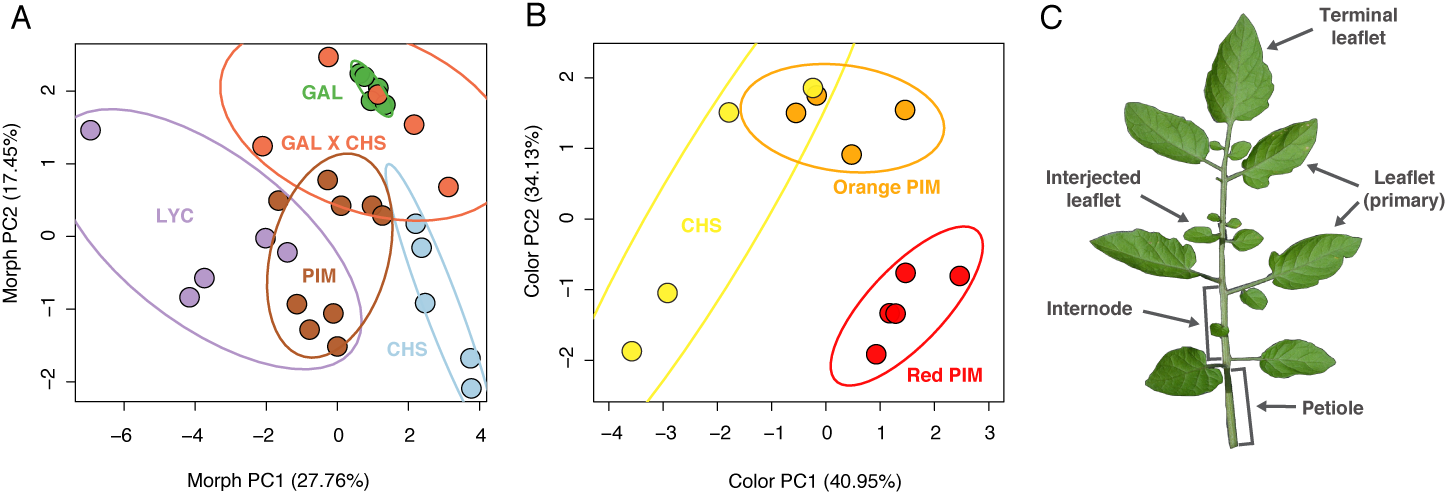
Quantitative morphological diversity of the Galápagos tomatoes. (A) First two principle component axes of morphological variation (leaves and fruit color) for five representative collections of each island taxon. (B) First two principle component axes of fruit color profiles (RGB & HSL) for collections at site MG115 and MG305. Some fruits were of low quality and were not included in color analyses. (C) Diagram of key morphological characteristics.

### 3.2 Santa Cruz

We revisited a total of 27 sites on Santa Cruz, 16 of which were previously documented to contain endemics or putative invasive-endemic hybrids (TGRC Passport data; Darwin, 2009; Nuez et al., 2004). We located only two endemic populations of CHS (MG115 & MG305) and no previously undocumented (new) populations. At site MG305 (Mina Granillo Negro), we observed a large mating group of Galápagos mockingbirds (*Mimus spp.*) carrying ripe CHS fruits. Mockingbirds have been previously hypothesized to be dispersers of wild tomato on the islands (Rick & Bowman, 1961; Nuez et al., 2004; Darwin, 2009), however systematic data to test this has not been collected and prior descriptions of frugivory have been limited to PIM. To our knowledge, we are the first to report observations of mockingbirds picking endemic tomatoes. The other relocated CHS population (MG115) was found 20m off Santa Cruz highway, at a site last documented by Nuez et al. (2004) as containing only CHS. However, we also observed one red- and two orange-fruited PIM individuals within 10m of endemic plants. While most revisited sites were proximal to human developments (i.e., in Puerto Ayora or along Santa Cruz Highway), we also visited five remote, uninhabited sites on the northern coasts of Santa Cruz and the adjacent island of Baltra (Figure 1). No tomato plants could be located at these sites. The absence of endemic populations was however most striking in Puerto Ayora—the largest city on Santa Cruz—and at the Charles Darwin Research Station (CDRS; Figure 1), where none of the six documented populations could be relocated. The most recent collection here—behind the entomology building at the CDRS—was made in 2002 by Sarah Darwin, but we found that this site had been recently cleared for an experiment involving cacti. These Puerto Ayora populations were absent in both 2018 and 2019.

Additional searches on Santa Cruz revealed multiple sites of PIM, LYC, and CER (Table 1). PIM was the most abundant, with 13 populations located; eight of these were polymorphic for fruit color, with approximately 30% of the plants in each population having orange instead of red fruits (Figure 3A & 3B). Orange-fruited CHS have been previously documented on Santa Cruz, and fruit color has been used as a diagnostic trait for endemic Galapagos species (Rick, 1971). However the plants identified here were identical to the coexisting red-fruited PIM plants in all other aspects except for fruit color (mean color PC1_Orange_ = -1.96, mean color PC1_Red_ = 1.57, P < 0.01; Table 3; Figure 2A & 2B), and also closely resemble PIM specimens from the islands as described in Darwin (2003). The only other notable difference detected between the red and orange biotypes was in the number of interjected leaflets; orange-fruited plants had on average two fewer interjected leaflets (P = 0.044; Table 3). For these reasons we refer to these biotypes as orange-fruited PIM (PIMO) and to these populations as polymorphic PIM, although we note that their status is being further evaluated with genetic data (see Discussion). Of the four locations where PIMO was observed, the new trash dump site (MG206) was previously reported to contain CHS in the surrounding mesic scrubland by Darwin (2000), but we found no endemic individuals at this location. In addition to these polymorphic populations, we located one population that appeared to be solely orange-fruited (MG113), and four that were solely red-fruited (Table 1). LYC and CER were found growing sporadically throughout the island, sometimes co-occurring with PIM (e.g., at MG118).

**Fig. 3.**
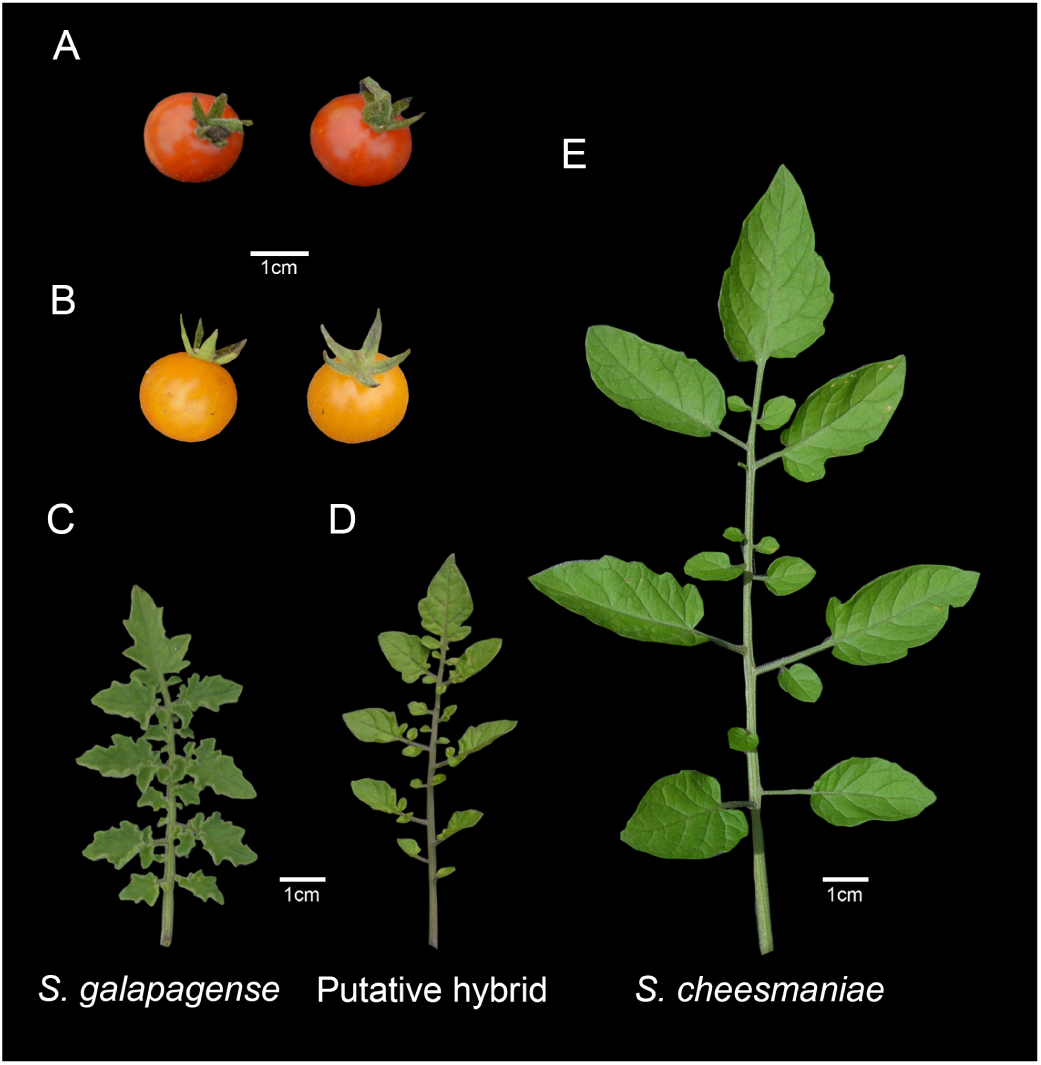
Qualitative morphological diversity of Galápagos tomatoes. (A & B) Red- and orange-fruited forms of *Solanum pimpinellifolium* from the polymorphic Mina Roja population on Santa Cruz. (C, D, & E) Representative leaves of *Solanum galapagense*, a putative CHS-GAL hybrid, and *Solanum cheesmaniae*, respectively, from the well (El Pozo) on Isabela.

### 3.3 Isabela

Isabela—the largest island in the archipelago, and the least developed of the three islands visited—contains 27 documented collection sites (TGRC passport data; Darwin, 2009; Nuez et al., 2004), however many of these are remote and inaccessible over land. Of the seven sites we revisited, all were in the Puerto Villamil and Santa Tomas regions on the South side of the island, and only two retained individuals of the endemic species. These two populations—one CHS and one GAL (MG120)—were growing sympatrically at a manmade well (El Pozo) adjacent to the Laguna de Manzanilla, a site first observed by Darwin (2002). Based on leaf morphology, several individuals collected at this site appeared to be hybrids between CHS and GAL (Figure 2A; Figure 3C, 3D, 3E). Putative hybrids had leaves of similar size to GAL, but leaflets that lacked secondary segments (i.e. leaflet margins were largely unlobed) like CHS; overall leaflet size was also more similar to GAL, but the orientation, distribution, and shape (i.e. the absence of tertiary lobing) of secondary and tertiary leaflets reflected CHS. In addition to this historical site, we also located two additional, previously undocumented, populations of GAL along the road to the highland region of Santa Tomás (MG124, MG127), less than 1 km from MG120. These individuals were similar in habitat and morphology to the well population.

Unlike on Santa Cruz, on Isabela we found PIM to be rare; CER was instead the major invasive type. Large (>500 plants) and previously undocumented populations of this variety were found at the Isabela trash dump (MG125) and in the highland town of Esperanza (MG126), a pattern consistent with the reported weedy and invasive distribution of this type throughout the tropical regions of South and Central America (Rick, 1973).

### 3.4 San Cristóbal

San Cristóbal contains only four historical collection records for endemics. Two of these are located in Puerto Baquerizo—the largest city on the island—and both were described as CHS (TGRC Passport data). Despite our searches, no endemic individuals could be located at these sites in either field season (Figure 1). Instead, LYC and PIM—which had been previously documented in Puerto Baquerizo (SCD901; Darwin, 2009)—were found within 0.5 km of the historical CHS collections (Table 1; Figure 1). East of Puerto Baquerizo—in the highland region of Progresso—contains no formal historical records of endemics or invasives. However, we explored this region and found two new populations of PIM (MG105 and MG107) as well as three cultivated LYC individuals (MG106).

## 4 Discussion

Our observations indicate that many historical populations of CHS and GAL on San Cristóbal, Santa Cruz, and Isabela have disappeared within the last 60 years, including several within no more than 16 years. Moreover, with the exception of the 1-2 novel GAL populations on Isabela, this loss of endemic populations does not appear to be balanced with the establishment of new endemic populations. For many sites our findings were consistent across two consecutive field seasons, suggesting that our observations are not solely due to transient yearly fluctuations in population presence and absence. Rather, they point to systemic shifts in species abundances on the islands, both for endemic and invasive species. The synergistic effects of expanding human developments—which began soon after initial collections were made in the 1950s and 60s (Quiroga, 2018)—along with the introduction of invasive species and interspecific hybridization, are potential drivers of this loss of endemic diversity.

### 4.1 Urbanization contributes to invasion and the loss of biodiversity

Urbanization has had numerous evolutionary and ecological consequences on the Galápagos (De Leon et al., 2018; Hendry et al., 2006; Sequeira et al., 2016; Tanner Perry, 2007), indicating that human-induced habitat disruption is one of the foremost threats to biodiversity on the islands (Pizzitutti et al., 2017; Quiroga, 2018). Our findings similarly suggest that urbanization has played an important role in the loss of endemic CHS and GAL populations. For all but five sites (North Santa Cruz & Baltra), endemic populations that could not be relocated were close to or within a human-disturbed habitat. The most striking instance was in Puerto Ayora, Santa Cruz where we were unable to locate endemic plants at any of the six collection sites visited. Such cases of local extinction can result in the irreversible loss of locally-unique trait variation, because genetic and phenotypic diversity is typically high between populations of endemic tomatoes in spite of low within-population variation due to high rates of selfing (Darwin, 2009; Pailles et al., 2017; Rick, 1983; Rick & Fobes, 1975). This loss is additionally troubling as, unlike many Galápagos plants, the endemic tomatoes have historically been an important source of traits for crop improvement (e.g., jointless pedicles in Puerto Ayora populations [Rick, 1967], high beta-carotene [Mackinney et al., 1954; Stommel, 1994], sugar content [Poysa, 1993], drought tolerance [Rush & Epstein, 1981], and insect resistance [Firdaus et al., 2013]; Rick, 1979; Nuez, 1995). In addition to local extinction, urbanization also appears to have facilitated the invasion of introduced species. At at least one site (MG206), invasives appear to have replaced previous endemics, and the observed abundance of invasive populations is associated with human population sizes on each island: on Santa Cruz (approx. 15,000 inhabitants) we observed 16 invasive populations, followed by 11 on San Cristóbal (approx. 7,000 inhabitants) and 5 on Isabela (approx. 2,000 inhabitants; Ecuadorian National Institute of Statistics and Censuses: http://www.ecuadorencifras.gob.ec). These patterns suggest that urbanization is one root cause of invasion, especially—as for many of the censused sites–in areas accessible by land and at sites adjacent to urban developments. Endemic populations on un-inhabited islands may be less prone to local extinction and/or interactions with invasives, especially if humans are the major vector of invasive tomato on the Galápagos. Nonetheless, we also describe the absence of five CHS populations in remote, undisturbed areas of Santa Cruz and Baltra, suggesting that the risk of extirpation could be high even in areas without invasives. The specific role of human introduction in invasive abundance and its subsequent effects on endemic diversity can be corroborated with additional censusing in more remote and/or undisturbed regions of the archipelago, especially on islands that have never hosted human settlers.

Recent anthropogenic changes could also act in concert with other non-anthropogenic factors. For example, many historical records of these species, and our observations here, indicate that population sizes are frequently small and suggest that intrinsic demographic fluctuations and local population turnover might be relatively rapid. Acute climatic changes could also exacerbate patterns of population turnover. For example, during our 2018 survey the islands were experiencing a three-month drought (which occurs periodically because of the El Niño Southern Oscillation) that could have contributed to the absence of endemic populations in coastal areas. However, we also did not relocate several endemic populations at higher elevations (Table 1) where it is typically moist year-round; in addition, extirpation was consistent at many sites across two consecutive years. Our observation of several missing remote populations on Santa Cruz and Baltra may point to more consistent climatic shifts and subsequent maladaptation as contributing to population loss. Censusing of these sites across multiple years and during different seasons/months will be necessary to evaluate the long-term effects of climate change on population persistence. Nonetheless, the disappearance of most historical populations, the clear association of this pattern with disturbed sites, and the lack of new (previously undocumented) endemic populations, all suggest that any natural population processes are being amplified by recent anthropogenic change. These potentially complex interactions between anthropogenic effects and natural demography highlight the need for more sustained population monitoring to better understand the nature and extent of conservation threats to these and similar species.

### 4.2 Human activity may promote endemic-endemic and invasive-endemic hybridization

Human activity can facilitate hybridization between species (Anderson & Stebbins, 1954; Todesco et al., 2016), either through the disturbance of native habitats that provides contact between isolated taxa (Anderson & Stebbins, 1954; Anderson, 1948; Simovich et al., 2013; van Hengstum et al., 2012) or through the introduction of non-native species (Fuller, Nico, & Williams, 1999; Maschinski, Sirkin, & Fant, 2010; Riley, Shaffer, Voss, & Fitzpatrick, 2003; Strong & Ayres, 2013). This appears especially true on the Galápagos where greater than 70% of alien plant species originate from human introductions (Quiroga, 2018), several of which have been implicated as potentially hybridizing with endemic congeners (Chaves, 2018; Torres & Gutierrez, 2018; Wendel & Percy, 1990).

Past observational and genetic evidence for hybridization between tomato species on the Galápagos is mixed—with evidence both for (Darwin, 2009; Pailles et al., 2017) and against (Nuez et al., 2004) intercrossing. Nonetheless, several biological factors suggest that the potential for gene flow between sympatric species is high. Intrinsic crossing barriers are known to be weak between all four taxa (Rick, 1956; Rick & Bowman, 1961; Rick & Fobes, 1975) and all co-occurring species were also co-flowering at the time of our survey. Pollinator observations for these species are limited, but movement of the polylectic Galápagos carpenter bee (*Xylocopa darwini*) between CHS and GAL has been documented on Isabela (Darwin, 2009). Movement between invasives and endemics is also expected to occur, as floral morphology is very similar among the four species (all flowers are yellow, rotate, and buzz pollinated).

The observation of individuals on Isabela that are intermediate for leaf shape/size between the sympatric CHS and GAL populations (Figure 2A; Figure 3C-E) suggests past or ongoing hybridization that may be influenced by human activity. Although perhaps not as troubling as intercrossing between an invasive and endemic, hybridization in this scenario could nonetheless result in the loss of genetic diversity and/or reduced genome integrity. The sympatric populations found at the well on Isabela are surrounded by a large gravel pit created to extract rock for the building of an airport in 1996 (Darwin et al., 2003). Collection records from prior years do not indicate populations at this location nor are the two species commonly found in sympatry (4/87 sites; TGRC passport data). Although endemic populations of intermediate morphology have been collected on Isabela as early as 1906 (Rick, 1963, 1971; Roger Chetelat pers. comm.), their ongoing co-occurrence at this location in 2018, first reported in 2000 (Darwin, 2009), and the possible hybridization between them is potentially the result of recent contact between two geographically isolated lineages because of habitat disruption.

Our observation of populations polymorphic for fruit color on Santa Cruz is also suggestive of hybridization, in this case between yellow- or orange-fruited endemics and introduced red-fruited PIM. The majority of collection records suggest that CHS typically has pale yellow fruits. However, orange-fruited CHS populations have been documented on Santa Cruz in Puerto Ayora that were initially classified as “Galápagos *S. pimpinellifolium*”; these populations share many characteristics in common with PIM but lack deep red fruits (TGRC passport data; Roger Chetelat pers. comm.). We cannot yet rule out the possibility that the orange-fruited PIM described here are in fact CHS. However, morphological comparisons to historical records suggest key differences present in our orange-fruited PIM, including their lack of lobing on primary and secondary leaflet margins and the presence of jointed pedicles. Such characteristics are more consistent with other red-fruited PIM collected on Santa Cruz (e.g., LA3123) than CHS. In contrast to the putative CHS-GAL hybrids on Isabela, these intermediate individuals were highly abundant yet phenotypically similar to PIM in all respects except for fruit color and number of interjected leaflets Table 3). The historical proximity of orange-fruited PIM-like CHS populations on Santa Cruz may help to explain how hybridization between species which typically show much more divergent traits could result in populations varying only in fruit color. Orange fruit color is extremely rare in mainland PIM (reported in 3 of 270 documented populations; TGRC Passport Data) and thus this shift in color has almost certainly evolved after colonization of the Galápagos —either as a result of hybridization or, alternatively, due to alleles that evolved in situ. Regardless, because both endemics also have lighter fruit colors, the observation of high frequencies of orange-fruited PIM-like individuals raises the intriguing possibility that selective conditions specific to the Galápagos have consistently favored shifts from red to lighter fruit colors. The source of this putative selection remains to be determined.

In general, our observational data generate several hypotheses that are testable with future field surveys and/or genetic analyses. First, how does proximity to human settlements explain the presence/abundance of invasive populations? Documentation of invasive tomato in remote areas of the archipelago is lacking, but will be important for evaluating the impact of anthropogenic activity on native species biodiversity. Second, are invasive and native species hybridizing? We observe several phenotypic patterns that may be consistent with interspecific hybridization. Distinguishing species admixture from intraspecific variation will ultimately require a genetic/genomic analysis of shared ancestry in populations displaying intermediate morphologies. Finally, what is the significance of orange fruit color, and what is its origin in PIM? The convergent evolution of orange fruits among invasive PIM and its native congeners suggests that specific environmental factors on the islands may select for lighter fruits. Lighter fruit color could have evolved in PIM either via introgression from endemic CHS or GAL or via a *de novo* transition. These alternative modes of convergence are distinguishable using genomic data (Lee & Coop, 2016), and the potential benefit of lighter fruits can be evaluated using *in situ* disperser color preference experiments.

## 5 Conclusion

Our field observation data suggests the occurrence of rapid population loss in the Galápagos tomatoes. This pattern appears to be driven primarily by the combined effects of expanding human developments, habitat disturbance, and the introduction of invasive relatives. Patterns of population loss in remote regions of the archipelago where human-induced disruption can be ruled out may also point to a role for sustained changes in the local climate as contributing to extirpation. Regardless of their causes, these findings raise questions about the current IUCN designations of “least concern” for CHS and GAL (Tye & Siemens, 2014). Further exploration of remote and/or undisturbed sites will be essential to determine whether these threats are experienced species-wide or generally localized to areas near human developments. However, for endemic populations on the three largest islands, our results are striking and highlight the fragility of these island species as well as the need for sustained population assessments in this and other understudied endemic plants.

## Acknowledgements

We thank the Galápagos Science Center staff on San Cristobal for logistic and permitting support and the Galápagos National Park for assistance locating and sampling endemic populations. Additional on-site field support was provided by Marcello Loyola and Genaro Garcia. Roger T. Chetelat, Iris Peralta, and two anonymous reviewers provided helpful comments that greatly improved the manuscript. This work was supported by a United States National Science Foundation award (IOS 1127059) to LCM. The authors declear no conflicts of interest. All field collections were made with appropriate permits and prior authorization by the Galápagos National Park and Ecuadorian Ministry of Environment.

